# Spatially distributed representation of taste quality in the gustatory insular cortex of awake behaving mice

**DOI:** 10.1101/2020.07.01.183095

**Authors:** Ke Chen, Joshua F. Kogan, Alfredo Fontanini

## Abstract

Visual, auditory and somatosensory cortices are topographically organized, with neurons responding to similar sensory features clustering in adjacent portions of the cortex. Such topography has not been observed in the piriform cortex, whose responses to odorants are sparsely distributed across the cortex. The spatial organization of taste responses in the gustatory insular cortex (GC) is currently debated, with conflicting evidence from anesthetized rodents pointing to alternative and mutually exclusive models. Here, we rely on calcium imaging to determine how taste and task-related variables are represented in the superficial layers of GC of alert, licking mice. Our data show that the various stimuli evoke sparse responses from a combination of broadly and narrowly tuned neurons. Analysis of the distribution of responses over multiple spatial scales demonstrates that taste representations are distributed across the cortex, with no sign of spatial clustering or topography. Altogether, data presented here support the idea that the representation of taste qualities in GC of alert mice is sparse and distributed, analogous to the representation of odorants in piriform cortex.

## INTRODUCTION

In primary sensory cortices, the spatial representation of sensory information can be either segregated or distributed. Experiments in somatosensory, visual and auditory cortices demonstrated a globally ordered topographic map, in which neurons representing similar sensory features are clustered together [1–6]. In contrast, odorants in the rodent piriform cortex evoke distributed patterns of activity with no spatial clustering [7, 8].

The spatial representation of taste in the gustatory insular cortex (GC) has been debated over the years. Some evidence supports the existence of a “gustotopic” map [9–11], which features dedicated “hot spots” of narrowly tuned neurons exclusively responding to individual taste qualities. According to this model no taste coding occurs outside of these hotspots [11]. Other imaging studies, however, describe largely overlapping regions of GC that respond to multiple taste qualities and report the existence of broadly tuned neurons, hence challenging the existence of a strict topographic organization [12–14].

Regardless of the fundamental differences between imaging studies of GC, they all share a common caveat: they were performed in anesthetized rodents. Thus, even if a topographic organization of taste exists in GC of anesthetized rodents, it is unclear whether and how it would persist during wakefulness considering that sensory processing is significantly affected by the state of the animal [15, 16]. This study is designed to assess the spatial organization of taste-related information in awake, behaving animals.

GC is located deep in the ventrolateral portion of the forebrain, making its access challenging for direct optical imaging in awake behaving rodents. Thus, we relied on implanted microprisms [17] to monitor neural activity in awake behaving mice with two-photon and widefield calcium imaging. We trained mice to perform a cued-taste paradigm, in which subjects actively licked a spout for gustatory stimuli delivered after a go cue. Using two-photon calcium imaging, we found sparse representations of cue, licking and taste stimuli in the superficial layers of GC. We observed overlapping representations for these three signals. Of the taste-responsive neurons, some were narrowly tuned, responding to a single taste, while others were tuned more broadly. Within local fields of two-photon images (450 x 450 μm), taste-responsive neurons were not spatially clustered. To study the organization of taste responses in GC at a larger spatial scale, we applied a widefield imaging approach (2 x 1.6 mm field of view). Analysis of responses confirmed that even at this large spatial scale, taste representations did not show any spatial clustering.

Altogether, data presented here support the idea that the representation of taste qualities in GC is sparse and distributed, analogous to the representation of odorants in piriform cortex.

## RESULTS

### Neural activity evoked by cue, licking and taste

Calcium imaging signals were obtained from the superficial layers of GC in 13 mice trained to perform a cued-taste paradigm (**Figure 1A,** see methods). Mice were first trained to lick a central spout following the offset a two second cue to obtain water, and then habituated for seven days to receive one out of five possible stimuli delivered pseudorandomly at each trial (sucrose [200 mM], NaCl [100 mM], citric acid [20 mM], quinine [1 mM] and water). After habituation, mice showed comparable duration of licking (n = 13, S: 3.6 ± 0.1 s; N: 3.5 ± 0.2 s; CA: 3.5 ± 0.2 s; Q: 3.5 ± 0.1 s; W: 3.5 ± 0.1 s; One-way ANOVA, F(4,60) = 0.07, p = 0.99, **Figure 1B-C**) and no significant difference in inter-lick interval (One-way ANOVA, F(4,60) = 0.04, p = 0.99). This similarity in licking behaviors was acquired with habituation (on day 1, licking responses differed according to palatability – see methods).

**Figure 1:**
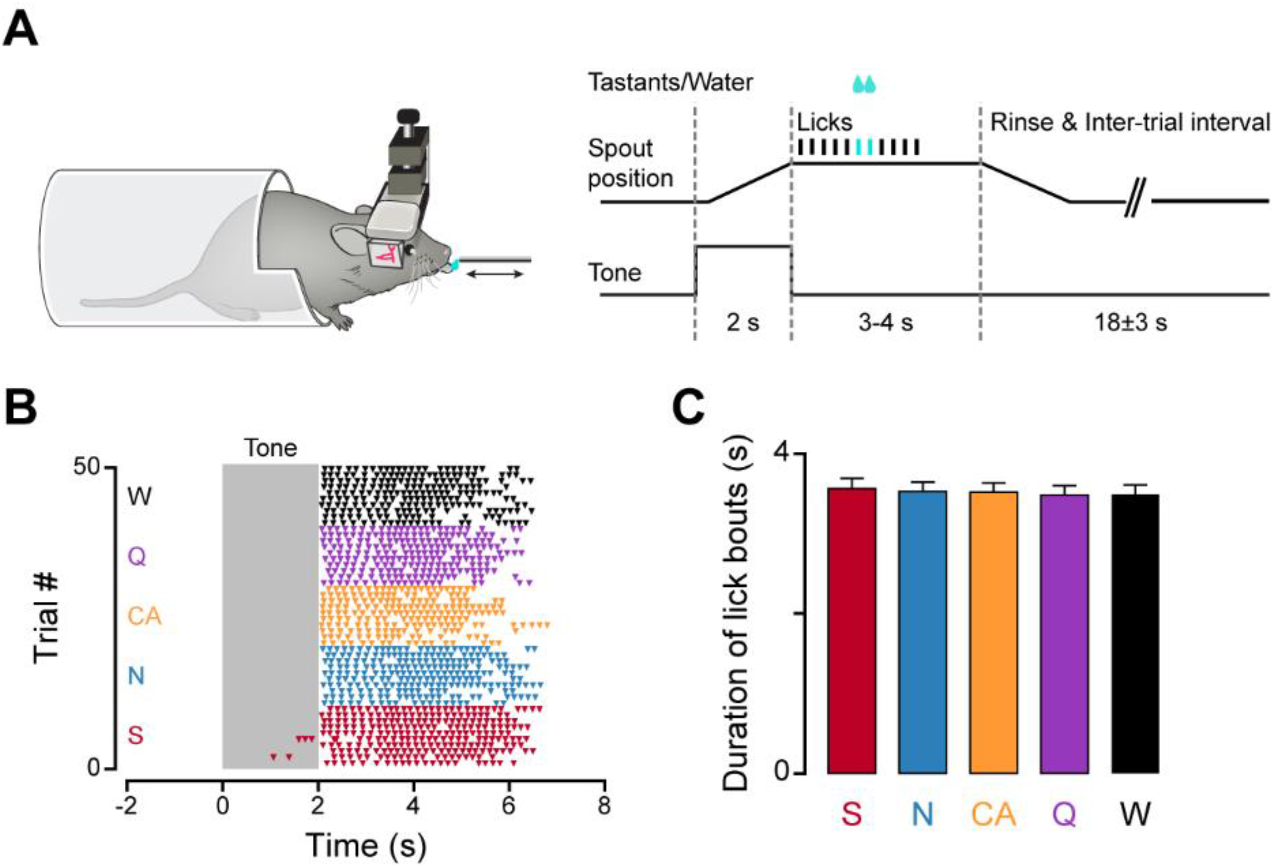
Behavioral paradigm. **A**, Left panel: Sketch showing a head-fixed, prism-implanted mouse licking a movable spout. Right panel: schematic diagram of the structure of each trial. **B,** Representative raster plot of licking to the five gustatory stimuli (sucrose: S, red; NaCl: N, blue; citric acid: CA, gold; quinine: Q, purple; water: W, black) in the cued-taste paradigm after habituating mice to the five tastants. Time 0 is the onset of the auditory cue and the shaded area represents the 2 s long auditory cue. Each triangle marker represents an individual lick. **C**, Bar plots representing the average duration of licking bouts (n = 13 mice) for the five gustatory stimuli after habituating mice to the cued-taste training. Error bars represent the standard error mean (SEM). One-way ANOVA with post hoc Tukey’s HSD test, p > 0.05.

To monitor calcium signals, we expressed the genetically encoded calcium indicator GCaMP6f [18] in GC (**Figure 2A**). To verify GCaMP6f expression in GC, in a subset of mice (n=3) we also injected an anterograde tracer (AAV1-CB7-CI-TurboRFP) into the taste thalamus (ventral posteromedial parvocellularis, VPMpc). As seen in **Figure 2A**, turboRFP-positive thalamic fibers (magenta) were colocalized with the neurons expressing GCaMP6f (green) in GC (**Figure 2A**).

**Figure 2:**
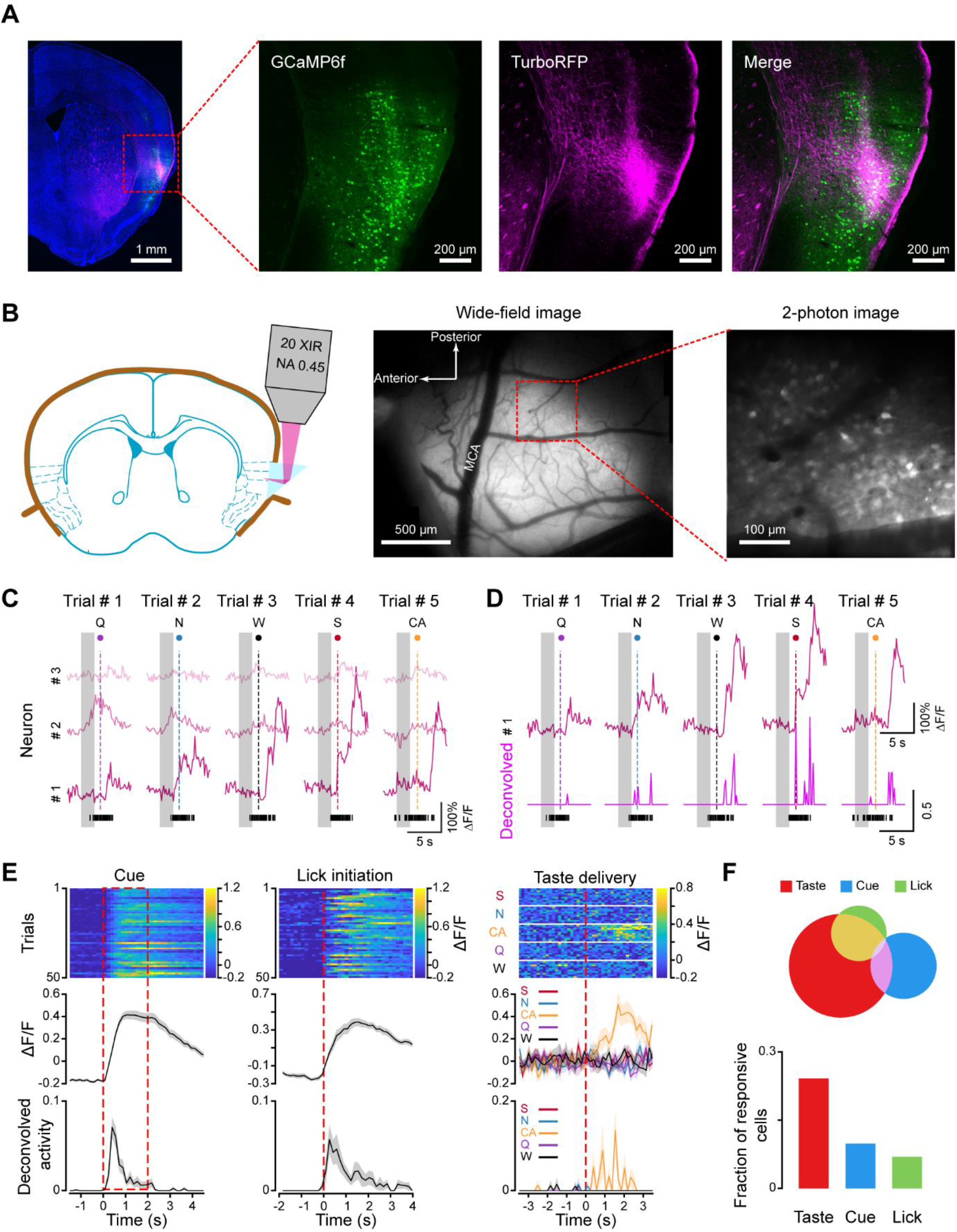
Two-photon calcium imaging of neural activity in GC of awake behaving mice. **A**, Histological images showing the expression of GCaMP6f (green) and anterograde TurboRFP-labeled VPMpc thalamic fibers (magenta) in GC. **B**, Left panel: schematic showing the prism positioned on the surface of GC with a 20 x air objective (NA: 0.45) used for imaging. Middle panel: a widefield image showing the expression of GCaMP6f (white) and the middle cerebral artery (MCA). Right panel: a two-photon image from the representative field marked on the widefield image. **C**, Representative calcium traces (ΔF/F) of three cells (# 1, # 2, # 3) in five consecutive trials. Gray bars represent the 2 s long cue. Dashed lines represent the delivery of the five gustatory stimuli (in the order of Q, N, W, S and CA). Black vertical ticks at the bottom represent licks. **D**, Calcium traces (ΔF/F) from cell # 1 in panel **C** and corresponding deconvolved neural activity (magenta traces) in five consecutive trials. Black vertical ticks at the bottom represent licks. **E**, Three representative neurons responding to cue (left), licking (middle) and tastants (right). Top panel: heatmap for the changes in fluorescent intensity (ΔF/F) evoked by cue (left), lick initiation (middle) and tastants (right). Each row represents a trial. Middle panel: average change of fluorescent intensity. Bottom panel: average deconvolved neural activity evoked by the cue, licking initiation and gustatory stimuli. For cue response, time 0 is the onset of the auditory cue. For licking response, time 0 is the initiation of licking. For taste response, time 0 is the delivery of the tastants. The shaded area around the curve indicates the SEM. **F**, Quantification of neurons responsive to cue, licking initiation and tastants. Top panel: Venn diagram showing the overlap of neurons representing cue, lick and tastants. Bottom panel: bar graph showing the fraction of neurons responsive to cue, lick and tastants.

To directly monitor neural activity from the superficial layers of GC (**Figure 2B**), we used two-photon calcium imaging. This approach allowed us to simultaneously record 50-150 neurons from mice engaged in the task (**Supplementary video 1**). We applied a constrained non-negative matrix factorization (CNMF) algorithm [19] to automatically segment regions of interest (ROIs, putative cells), extract calcium traces and deconvolved activity of each cell (**Figure 2C**). Deconvolution allowed us to disambiguate responses to cue, licking initiation and tastants (**Figure 2D**). In total, we recorded 1137 neurons from 10 mice (16 sessions). Consistent with previous studies [20–23], neurons in GC responded to all the events in the task: anticipatory cue, licking initiation and gustatory stimuli (**Figure 2E**). In total, we observed 9.9% (112/1137) of neurons responded to cue, 6.9% (79/1137) of neurons responded to licking initiation and 24.2% (275/1137) of neurons responded to the gustatory stimuli (**Figure 2F**). We also observed cells with overlapping responses, 24.1% (27/112) of cue-responsive and 77.2% (61/79) of lick-responsive neurons also responded to gustatory stimuli. Overall, the responses to the cue, lick initiation and taste in the superficial layers of GC were sparse.

Next, we analyzed responses to each of the five stimuli. Taste-responsive neurons responded to S, N, CA, Q and W in the following proportions: S: 40.7% (112/275), N: 40.4% (111/275), CA: 37.1% (102/275), Q: 39.6% (109/275) and W: 43.6% (120/275) (**Figure 3A-B**). No significant difference was observed in the average evoked response to the five tastants (for evoked ΔF/F, S: 0.27 ± 0.01, N: 0.32 ± 0.02, CA: 0.28 ± 0.02, Q: 0.27 ± 0.02, W: 0.28 ± 0.02, one-way ANOVA, , F(4,549) = 1.41, p = 0.23; for evoked deconvolved activity, S: 0.051 ± 0.003, N: 0.053 ± 0.003;CA: 0.051 ± 0.003; Q: 0.047 ± 0.003, W: 0.051 ± 0.003, one-way ANOVA, ,F(4,549) = 0.5, p = 0.74, **Figure 3C**). The fraction of taste-responsive neurons with best responses to the five tastants was also comparable (S: 20.4% [56/275], N: 17.4% [48/275], CA: 20.4% [56/275], Q: 19.6% [54/275], W: 22.2% [61/275], Pearson’s χ^2^ test, χ^2^ = 2.0, p = 0.74, **Figure 3D**). Based on evoked responses, we observed both narrowly and broadly tuned neurons, 47.6% (131/275) of neurons responded to only one tastant, 52.4% (144/275) responded to multiple tastants (**Figure 3E**). To further evaluate tuning, we applied a hierarchical clustering analysis to classify the taste responses. This analysis identified 16 clusters (**Figure 3F**). For each cluster, we calculated the entropy – a well-established measure for the breadth of tuning [24]. Five clusters had low entropy (0.019 ± 0.01), representing neurons (50.2% [138/275]) that were narrowly tuned to the five tastants. The other 11 clusters had high entropy (0.62 ± 0.04), representing neurons (49.8% [137/275]) broadly tuned to multiple tastants (**Figure 3G**). Thus, we found a range of both broadly and narrowly tuned taste-responsive cells in superficial layers of GC, consistent with previous imaging studies in anesthetized mice [13].

**Figure 3:**
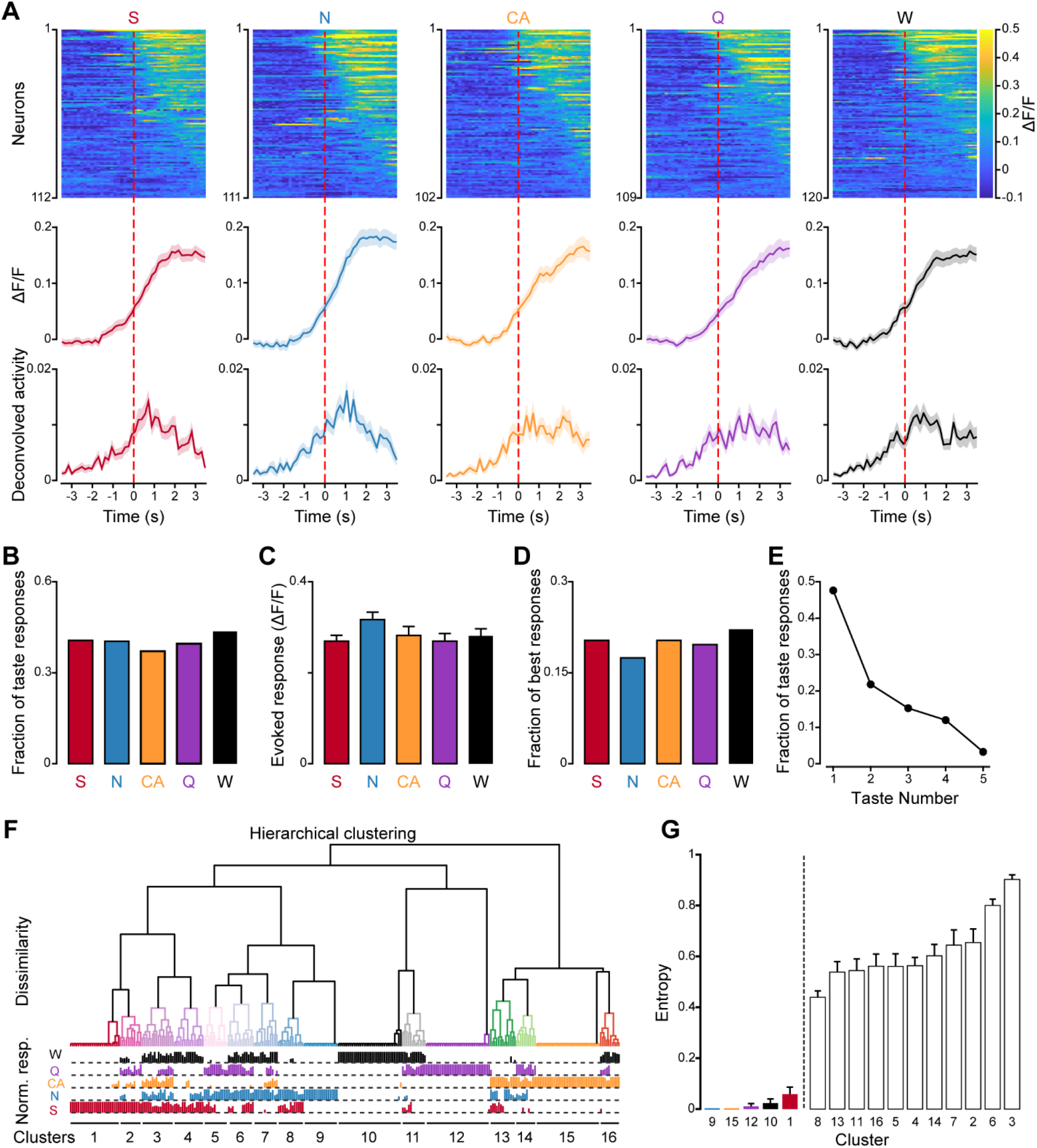
Taste response in GC from awake behaving mice. **A**, Top panel, heatmap of population activity for each of the five gustatory stimuli. Each row represents a single neuron. The color represents the change of fluorescent intensity (ΔF/F). Middle panel, average of the change in fluorescent intensity (ΔF/F) across neurons responsive to each of the five gustatory stimuli. Bottom panel, average of the deconvolved activity across neurons responsive to each of the five gustatory stimuli. The dash line (time 0) represents the taste delivery. The shaded area around the curve indicates the SEM. **B**, Bar graph showing the fraction of taste-responsive neurons to each of the five tastants. **C**. Bar graph showing the average amplitude of evoked responses to the five tastants. **D**, Bar graph showing the fraction of neurons with best responses to each of the five tastants. **E**, Tuning curve showing the fraction of neurons responding to 1, 2, 3, 4 and all 5 tastants. **F**, Top panel: dendrogram of hierarchical clustering analysis based on normalized responses. The 16 colors of the dendrogram represent 16 clusters. Bottom panel: bar graph of normalized responses. Each column represents an individual neuron. Each row represents the normalized response to each tastant. **G**, Bar graph showing the average entropy of cells belonging to each cluster.

### Spatial representation of taste quality

Visual inspection of taste responses suggests the absence of any spatial clustering of taste-responsive neurons for any of the five stimuli (S: sweet, N: salty, CA: sour, Q: bitter, W: water). **Figure 4A** shows a representative two-photon imaging field with neurons responding to the five stimuli. To quantify the spatial distribution of taste responses, for each stimulus we compared the pairwise distance between taste-responsive neurons with a null distribution (see method, **Figure 4B**). We used a significance threshold of 0.05 to avoid an excessively stringent criterion for identifying clustering (see methods for results for the stricter 0.01 threshold). We observed that in the majority of individual sessions (S:13/16 sessions, N: 15/16 sessions, CA: 15/16 sessions; Q: 15/16 sessions; W: 14/16 sessions), the distance between taste-responsive neurons was not significantly smaller than the distance between randomly selected neurons, suggesting that there was no spatial clustering. In the instances with significant smaller pairwise distance between taste-responsive neurons (n = 5 sessions), the average intra-cluster distance between neurons (see method, **Supplementary Figure 1B**) was not significantly different from inter-cluster distance, indicating neurons responding to each quality were still intermingled (**Supplementary Figure 1**). Furthermore, in these instances we observed that 54% (41/76) of taste responsive neurons were broadly tuned, a result incompatible the idea that these responses may represent some remnants of hot spots. In addition, we did not observe consistent spatial clustering for neurons responsive to the anticipatory cue and licking (**Supplementary Figure 2**).

**Figure 4:**
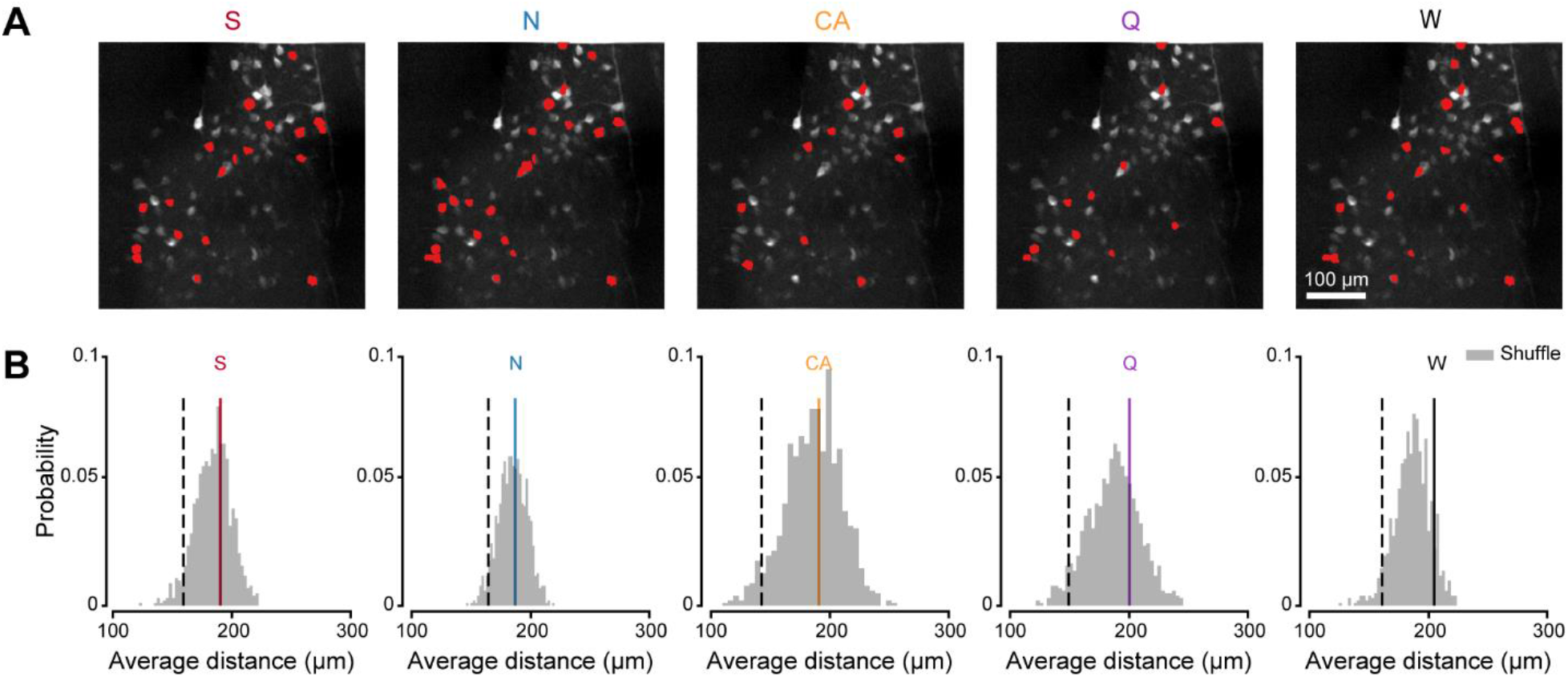
Spatial representation of taste qualities with two-photon imaging. **A,** Representative two-photon images from the same field showing the location of neurons (red markers) responding to S, N, CA, Q and W. **B** Representative histograms showing the distribution of average pairwise distance between randomly chosen neurons (grey). The black dashed lines mark the boundary of the lowest 5% average pairwise distance for the random distribution. Colored lines represent the average pairwise distance between neurons with response to S (red), N (blue), CA (orange), Q (purple) and W (black), as shown in **A**.

One of the potential limitations of our two-photon approach is a field of view constrained to 450 x 450 μm. To evaluate the spatial organization of a larger portion of GC, we applied a widefield imaging approach, which provided a ~2 x 1.6 mm field of view with single-cell resolution (**Figure 5A, Supplementary Video 2**). We used a CNMF-E algorithm, an extension of the CNMF for one-photon imaging, to automatically extract the location of ROIs (putative cells), calcium traces and deconvolved activity for each cell (**Figure 5A**). In total, we recorded 3325 putative neurons from 4 mice (including one mouse recorded with two-photon imaging before) with widefield imaging. The data confirmed the sparseness of responses to cue (6.32% [210/3325]), lick initiation (5.71% [190/3325]) and taste (18.59% [618/3325]), already observed with two-photon imaging. The prevalence of cue and taste responses observed with widefield imaging was significantly lower than that seen with two-photon imaging (cue responses: Pearson’s χ^2^ test, χ^2^(1) = 15.8, p < 0.001; lick responses: Pearson’s χ^2^ test, χ^2^(1) = 2.1, p = 0.15; taste responses: Pearson’s χ^2^ test, χ^2^_(1)_ = 16.2, p < 0.001). This difference may be related to widefield imaging being able to resolve activity only from more superficial layers compared to two-photon.

**Figure 5:**
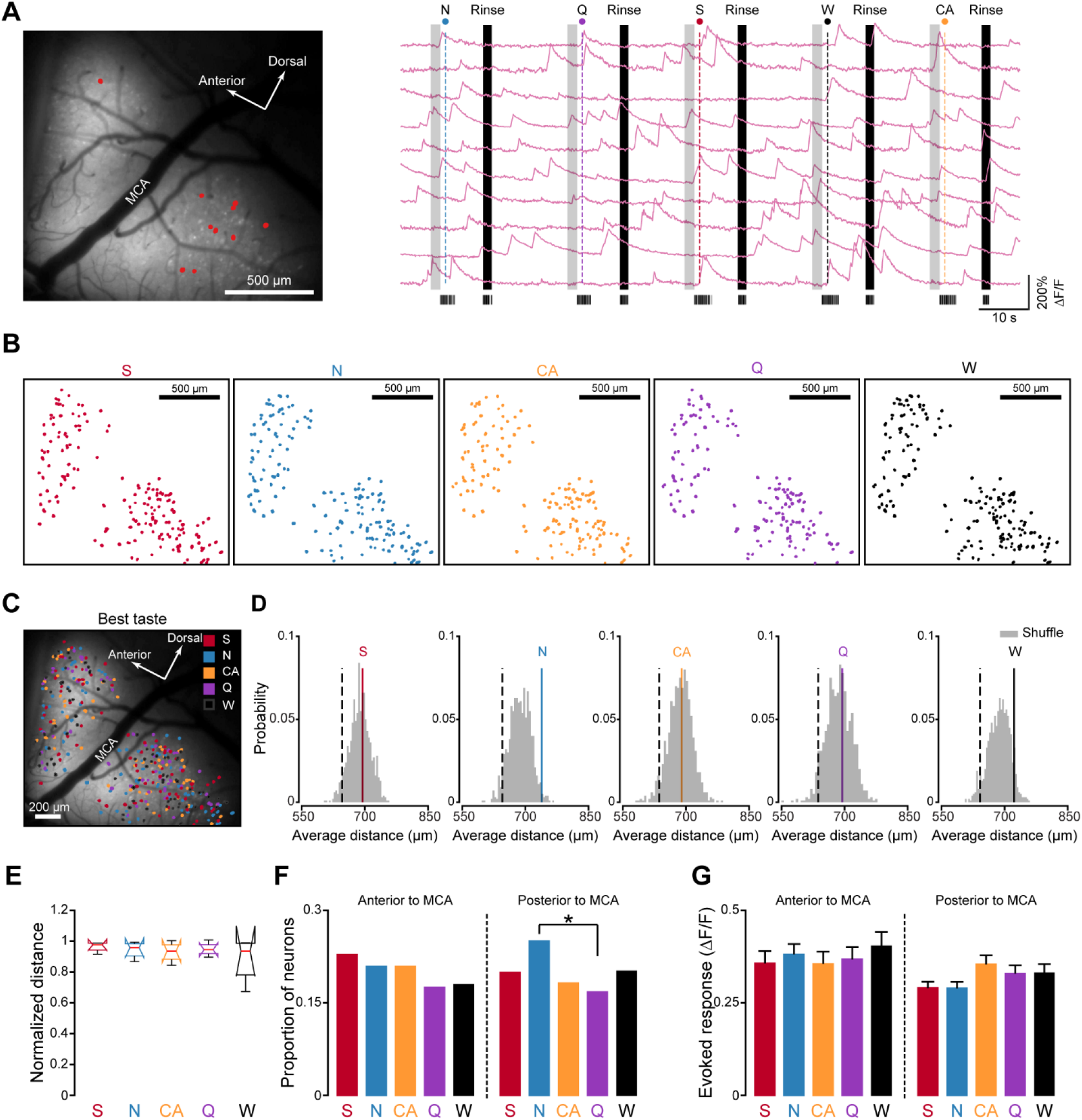
Spatial representation of taste qualities with widefield imaging. **A,** Left panel, a representative widefield image showing the expression of GCaMP6f (white) and the middle cerebral artery (MCA). Right panel, representative calcium traces (ΔF/F) of 10 example neurons (marked as red dots on the left widefield image) in 5 consecutive trials. Gray bars represent the 2 s auditory cue. Dash lines represent taste delivery. Black bars represent the rinse. The bottom black ticks represent licks. **B,** Representative spatial locations of neurons responding to S, N, CA, Q and W. The field of the spatial map is the same as the one shown in **A**. Notice the distributed organization of neurons responding to the five tastants. **C**, Representative spatial map of neurons with best responses to S (red), N (blue), CA (orange), Q (purple) and W (black). **D**, Representative histograms showing the distribution of average pairwise distance between randomly chosen neurons (grey). The black dashed lines mark the boundary of the lowest 5% average pairwise distance for the random distribution. Colored lines represent the average pairwise distance between neurons with best response to S (red), N (blue), CA (orange), Q (purple) and W (black), as shown in **C**. **E**, Box plot of the normalized intra-cluster distance of neurons with best responses to S, N, CA, Q and W. Distance was normalized to the average inter-cluster distance (n = 4 sessions). **F**, Bar graph showing the proportion of taste-responsive neurons anterior to MCA (n = 206, left side of the vertical dash line) and posterior to MCA (n = 412, right side of the vertical dash line) with best responses to S, N, CA, Q and W. Pearson’s χ^2^ test with Bonferroni correction, * represents adjusted p < 0.05. **G**. Bar graph showing average amplitude of evoked best responses to the five tastants for neurons anterior (n = 206, left side of the vertical dash line) and posterior to MCA (n = 412, right side of the vertical dash line). Error bars represent the SEM.

To assess the spatial organization of taste qualities at this larger field of view, we first calculated the pairwise distance between neurons evoked by each of the five tastants (S, N, CA, Q, W) as described above. As for the two-photon analysis, also in this case we used a significance threshold of 0.05 (see methods for results with a more stringent, 0.01, threshold). **Figure 5B** shows an example of spatial locations of neurons responding to the five stimuli in one imaging field. Taste-evoked responses for each gustatory stimulus were not clustered or segregated, but instead distributed across the field. Indeed, in the majority of imaging fields (S:3 of 4 sessions, N: 3 of 4 sessions, CA: 3 of 4 sessions, Q: 4 of 4 sessions, W: 3 of 4 sessions), distance between taste-responsive neurons was not significantly smaller than the distance between randomly selected neurons, suggesting that even at a larger field of view, responses to taste in the superficial layer of GC are randomly distributed. In the session (n =1) with significantly smaller pairwise distance between taste-responsive neurons, 59% of the neurons (33/56) were broadly tuned and the intra-cluster distance was not significantly different from inter-cluster distance, indicating that neurons responding to each quality were still intermingled (**Supplementary Figure 3A-C**).

This distributed organization of taste responses can be related to the relatively large proportion of broadly tuned neurons and to the duplication of those neurons in the analysis for multiple tastants. To adopt a more stringent criterion, we re-calculated the distance between neurons focusing exclusively on the best response to each of five stimuli (**Figure 5C-5D, Supplementary Figure 3D-E**). Visual inspection of the spatial map of best responses (**Figure 5C**), suggests that even in this scenario, responses to tastants were distributed. Indeed, in the majority of imaging fields (S:4 of 4 sessions, N: 3 of 4 sessions, CA: 4 of 4 sessions, Q: 4 of 4 sessions, W: 4 of 4 sessions), distance between neurons with best responses to each tastant was not significantly smaller than the distance between randomly chosen neurons. **Figure 5E** shows the normalized intra-cluster distance of neurons with best response to the five gustatory stimuli in 4 imaging sessions.

It has been previously reported that GC neurons anterior to the middle cerebral artery (MCA) respond more to sucrose and NaCl [11], and GC neurons posterior to MCA respond more to quinine [11, 25]. In our imaging dataset, we did not observe this trend. Indeed, a comparable proportion of GC neurons anterior to MCA responded to each of the 4 taste qualities (S: 22.8% [47/206], N: 20.9% [43/206], CA: 20.9% [43/206], Q: 17.5% [36/206], W: 18.0% [37/206], Pearson’s χ^2^ test, χ^2^_(4)_ = 2.57, p = 0.63, **Figure 5F**). GC neurons posterior to MCA showed a similar tendency of responding to the 4 taste qualities (S: 20.0% [82/412], 25.0% [103/412], 18.2% [75/412], 16.8% [69/412], 20.2% [83/412]), with a slightly higher tendency of responding to NaCl compared to quinine (Pearson’s χ^2^ test with Bonferroni correction, adjusted p = 0.04). Similar results were obtained when we focused exclusively on the amplitude of evoked best responses. No significant taste preference was found for neurons located anterior and posterior to MCA (anterior to MCA, evoked ΔF/F, S: 0.35 ± 0.03, N: 0.38 ± 0.03, CA: 0.35 ± 0.03, Q: 0.37 ± 0.03, W: 0.40 ± 0.04, One-way ANOVA, F(4,201) = 0.36, p = 0.84; posterior to MCA, evoked ΔF/F, S: 0.29 ± 0.02, N: 0.29 ± 0.02, CA: 0.35 ± 0.02, Q: 0.33 ± 0.02, W: 0.33± 0.02, One-way ANOVA, F(4,407) = 2.15, p = 0.07, **Figure 5G**).

In summary, our data show that in the superficial layers of GC of alert mice, taste is represented through sparse and distributed patterns of activity.

## DISCUSSION

This study evaluated the spatial representations of cue, licking, and taste qualities in the superficial layers of GC in awake behaving rodents. We trained mice in a cued-taste paradigm to lick a spout after an auditory cue to receive tastants. To monitor neural activity, we chronically implanted microprisms above the surface of GC and performed two-photon calcium imaging. We observed that the activity evoked by the cue, licking and all the gustatory stimuli was sparse. Analysis of taste-evoked responses showed a combination of narrowly and broadly tuned neurons. Taste representations were spatially distributed. To exclude that the lack of clusters was not related to the limited field of view of two-photon imaging (450 x 450 μm), we performed widefield imaging with cellular resolution over a 2 x 1.6 mm region of GC. Like with two-photon imaging, we found that GC neurons representing chemosensory information were largely scattered throughout the field and did not form isolated and selective clusters. These results support a spatially distributed coding scheme for taste-related information in the superficial layers of GC, analogous to the coding of odorants in the piriform cortex [7, 8].

### Multimodal responses in GC

The role of GC in processing taste-related information has traditionally been studied with electrophysiological recordings. Unlike the electrophysiological approaches used in GC, imaging allows for monitoring of large neural ensembles while preserving spatial information. Here, we imaged neural activity in the superficial layers of GC using two-photon and widefield calcium imaging. Consistent with electrophysiological studies [20–23, 26], we observed GC responding to taste qualities, cue, and licking, with a mixture of narrowly and broadly tuned neurons. Neurons could either respond exclusively to a single modality or to multiple modalities (i.e., a convergent representation of cue, licking and taste qualities). This observation re-affirms that GC is multimodal, and capable of encoding non-chemosensory, taste-related variables.

Though largely consistent, the observed responses to taste qualities (19-25%), cue (6-10%), and licking (~5%) were relatively sparse compared with previous work [20, 27, 28]. This discrepancy could arise from several factors. First, our study only examined excitatory responses, inhibitory responses were not included due to the difficulty of analyzing them in imaging datasets. Inhibitory modulation has been observed in GC during active licking [27]. Thus, by excluding inhibitory responses, our study underestimates responsiveness to taste, cue and licking. Second, our microprism-based imaging approach only allows for recording from neurons located in the superficial layers of GC, electrophysiological recordings generally sample from deeper layers. In sensory cortices, neurons in different layers can have varied tuning and response profiles, with some reports showing sparser representations of stimuli in superficial layers [29–31]. While our result is consistent with work on other sensory cortices, the representation of taste-related information across different layers of GC requires further study. Finally, the lower sensitivity of calcium imaging relative to single unit extracellular recordings may have played a role in underestimating neural responses.

In our dataset, the majority of GC neurons (60-70%) did not respond to taste, cue or licking. Neurons in GC have been shown to encode a broad range of cross-modal information, including olfactory, visual and somatosensory stimuli [20]. A subset of the non-responsive neurons may be involved in encoding this information and participate in the perception of flavor [32, 33] and the formation of associative representations triggered by anticipatory cues [20, 21, 26]. Moreover, GC neurons have been shown to encode cognitive variables associated with decision making [27, 34]. Hence, a proportion of the non-responsive neurons may also participate in encoding cognitive variables associated with the task. Future imaging of GC will require the use of more complex tasks that involve learning and decision making.

### Spatial representation of taste quality in GC

In sensory cortices, the spatial representation of sensory information can be either clustered or distributed. In primary somatosensory, visual and auditory cortices, neuronal responses are organized into a topographic map, with neurons encoding similar stimulus features, such as spatial proximity, orientation or frequency, clustering near each other [2, 3, 5, 6]. In contrast, the representation of olfactory information in rodent piriform cortex is sparse and distributed [7, 8].

In the past decade, several attempts have been made at applying optical imaging to study the spatial coding of taste quality in GC, and the results are discordant [11–14, 35]. Some studies describe spatial clusters tuned exclusively to individual stimuli with virtually no broadly tuned neurons inside or outside of the clusters [11]. Others find a combination of narrowly and broadly tuned neurons with no spatial clustering [13]. Regardless, all these studies have been conducted in anesthetized rodents. Sensory coding has been shown to be sensitive to anesthesia, thus it is unclear how these findings would extend to alert animals [15]. Here, we attempt to resolve some of these controversies by recording taste responses in GC of awake, behaving rodents. Using two-photon and widefield calcium imaging, we observed both narrowly and broadly tuned taste-responsive neurons that are spatially distributed throughout the superficial layer of GC.

Our experiments relied on single concentrations of each gustatory stimulus and hence caution should be taken in generalizing our results to all stimulus intensities. It is theoretically possible that spatial clustering may emerges only for selected stimulus intensities [36]. However, the concentrations adopted for our experiments are consistent with those widely used in the field [28, 37, 38] and for three stimuli (sucrose, citric acid and NaCl), we chose the same concentrations used in the study that described a strict topographic organization. [11]. For quinine we relied on a lower concentration than the aforementioned study (1mM vs 10mM) because 10mM is highly aversive and not suitable for an active licking paradigm. It is unlikely that high stimulus intensity may lead to spatially localized responses, as studies in gustatory sensory ganglion neurons demonstrate that high stimulus intensities increase, instead of reducing, the breadth of tuning [37].

While stimulus intensity is unlikely to account for the discrepancy between our findings and those reported in Chen et al, fundamental differences in experimental design may have played a key role. First, we used GCaMP6f, a more sensitive calcium indicator than bulk-loaded dyes. Second, taste responses were recorded from awake, behaving mice rather than anesthetized animals. Third, the method of taste delivery differed dramatically. In our experiments, mice received 2 drops of tastants by actively licking. In Chen et al., tastants were perfused into the oral cavity for 10 s. These factors may account for our different observations.

It is worth emphasizing that our results are consistent with other studies using intrinsic and two-photon imaging in anesthetized rodents which showed that regions in GC responding to different tastes are largely overlapping [12–14].

The studies relying on two-photon imaging, like ours, covered relatively small fields of view (450 x 450 μm). Thus, it could be argued that a spatial organization of taste responses might still exist on a larger scale, especially at the rostral and caudal extremes of GC. To address this concern, we performed widefield imaging with cellular resolution and imaged neural activity in GC at a large scale (2 x 1.6 mm). With this technique we still observed that responses evoked by taste qualities were distributed across the surface of GC. Even when we re-categorized neurons by their best responses – a procedure that could bias the analysis toward a topographic organization - representation of taste quality was spatially distributed.

Despite an overall distributed representation, there could still be a gradient of best responses in the anterior and posterior portions of GC. We separately looked at responses in regions anterior and posterior to the middle cerebral artery, a landmark that bisects GC, and still found little to no evidence for the spatial biasing of taste responses. These observations are consistent with electrophysiological studies in alert rodents showing that taste tuning does not depend on spatial location within GC [27, 28, 38]. Indeed, neurons in anterior or posterior GC show comparable tuning and tendency of responses to each taste quality [28].

Altogether, our data provide compelling evidence for a distributed organization of taste representations in GC, reminiscent of odorant coding in piriform cortex [7, 8]. This similarity suggests that chemo-sensation shares a distributed coding scheme differing from the topographical organization of visual, somatosensory and auditory systems.

## ACKNOWLEDGEMENTS

The authors would like to acknowledge Dr. Craig Evinger, Dr. Arianna Maffei, Dr. Joshua L. Plotkin, Dr. Daniel B. Polley, past and present members of the Fontanini and Maffei’s laboratories at Stony Brook University for their feedback and insightful comments. This work has been supported by National Institute on Deafness and Other Communication Disorder Grants R01DC018227 and R21DC017681 to AF.

## AUTHOR CONTRIBUTION

K.C., and A.F. carried out study conceptualization and experimental design. K.C. and J.K. performed calcium imaging, behavioral experiments and data analysis. All the authors contributed to writing the manuscript.

## MATERIAL AND METHODS

### Experimental subjects

Adult male mice (C57BL/6J, 12-20 weeks old, The Jackson Laboratory) were used for this study. We used exclusively male mice to reduce the possible variability associated with estrous cycle. Mice were group housed and maintained on a 12 h light/dark cycle with *ad libitum* access to food and water unless otherwise specified. All experimental protocols were approved by the Institutional Animal Care and Use Committee at Stony Brook University, and complied with university, state, and federal regulations on the care and use of laboratory animals.

### Surgical procedures for viral injection and prism implantation

Mice were anesthetized with an intraperitoneal injection of a mixture of dexmedetomidine (1 mg/kg) and ketamine (70 mg/kg). The depth of anesthesia was assessed by testing pinch reflex. Once fully anesthetized, mice were placed on a heating pad (DC temperature control system, FHC, Bowdoin, ME) to maintain the body temperature at 35 °C. The animal’s head was shaved, cleaned, disinfected (three alternating washes of iodine and ethanol) and fixed to a surgical stereotaxic apparatus. Carprofen (5 mg/kg) was injected subcutaneously for analgesia. Ophthalmic ointment was placed on eyes to prevent dehydration. The scalp was carefully cut open and the skull was leveled. A small craniotomy was drilled on the dorsal portion of the skull above left GC (AP: 1.2 mm, ML: 3.5-4.0 mm relative to bregma). A pulled glass pipette front-loaded with virus carrying GCaMP6f (AAV1-hSyn-GCaMP6f-WPRE-SV40, 2.3 × 10^13^ gc/mL, catalog # 100837-AAV1, Addgene) was lowered into GC (1.9-2.0 mm below the dura) and a microinjection syringe pump (UMP3T-1, World Precision Instruments) was used to inject a total of 200 nL virus at 1 nL/s. Two viral injections (100 nL each) were performed at two different anterior-posterior locations (1.2 mm and 0.9 mm anterior to bregma). After each injection, the pipette was left in place for five minutes before being slowly retracted. In a subset of mice (n= 3), we also injected 100 nL anterograde viral tracer (AAV1-CB7-CI-TurboRFP-WPRE-RBG, 2.2 × 10^12^ gc/mL, catalog # 105546-AAV1, Addgene) into the ventral posteromedial parvocellularis (VPMpc) of thalamus (AP: −1.8 mm, ML: 0.6 mm relative to bregma, DV: −4.0 mm below dura). The craniotomy was covered with silicone gel and the scalp was sutured close. After the surgery was complete, Antisedan (atipamezole hydrochloride, 1 mg/kg) and lactated ringer’s solution were administered subcutaneously to reverse anesthesia and for hydration respectively.

Two to three weeks after viral injection, mice were implanted with prisms. Mice were anesthetized with an intraperitoneal injection of a mixture of dexmedetomidine and ketamine as described above. Once fully anesthetized, mice were subcutaneously injected with carprofen (5 mg/kg) and dexamethasone (2 mg/kg). The left eye was sutured close. The scalp and the skin between the left eye and ear were removed. Bupivacaine (2.5 mg/mL, 0.01-0.02 mL) was injected into the temporalis muscles for local anesthesia. Portions of the temporalis muscles were removed, and a ~ 2.2 x 2.2 mm cranial window was opened on the lateral portion of the skull to directly expose the surface of GC (bottom of the craniotomy window was at the squamosal plate). The middle cerebral artery and rhinal vein were used as surgical landmarks for GC. A glass prism assembly was implanted to cover the craniotomy and secured in place with Vetbond and black dental acrylic. The glass prism assembly was fabricated by gluing with optic glue (NOA61) a coverslip (#1 thickness) onto the surface of a 2 mm prism (MPCH-2.0, Tower Optics) that faces the gustatory cortex and gluing another coverslip on the hypotenuse. A customized headpost was cemented to the dorsal portion of the skull for head restraint. Mice were injected with carprofen (5 mg/kg, subcutaneous) daily for three days after the surgery to reduce inflammation.

### Cued-taste paradigm

Following recovery, mice were placed on water restriction, with 1.5 mL water given daily for one week before training. Weight was monitored and maintained at > 80% of the initial weight before water restriction. In the first phase of training, mice were habituated to licking a spout after a 2 s cue to receive water. For each trial, a 2 s auditory tone was presented (2k Hz, 70 dB) and a motorized spout (X-LSM motor, Zaber) moved in front of the animal’s mouth. The offset of the tone and the end of spout movement were aligned. Mice were required to lick the dry spout once to trigger the delivery of two drops of water (3 μL each) and the spout remained in place for 3 s to allow the mouse to consume the water before retracting. A rinse with two drops of water was also introduced after each trial (a rinsed was introduced 7.8 ± 0.5 s after taste delivery). The inter-trial interval was 12 ± 2.5 s and an additional 25 s timeout was triggered if mice failed to lick after the cue. Once mice started to reliably perform a dry lick for water (1-2 sessions, performance > 90%), the number of dry licks required to trigger water delivery was increased (from 1 to 5). Mice were then habituated to the 5 dry licks (4-5 sessions). After water habituation training, training began with the five gustatory stimuli (sucrose [200 mM], NaCl [100 mM], citric acid [20 mM], quinine [1 mM] and water). Trials for each tastant were presented in a random order (random permutation, on average 20 trials for each taste). On day one, mice showed significantly longer duration of licking to S and N, than to CA and Q (S: 3.6 ± 0.1 s; N: 3.5 ± 0.1 s; CA: 2.9 ± 0.1 s; Q: 2.5 ± 0.2 s; W: 3.1 ± 0.1 s; One-way ANOVA, F(4,60) = 12.5, p < 0.001, post hoc Tukey’s HSD test, p <0.05). After a week of habituation, mice showed comparable duration of licking to each of the tastants, and imaging experiments started. Gustatory stimuli were delivered via a gravity-based taste delivery system. The spout was composed of five independent polyamide tubes, each connected to a taste line. An infrared beam (940 nm, powered by a fiber-coupled LED, Thorlabs) was positioned in front of the mouth for lick detection. Behavioral events and licking data were recorded with RHD2000 recording system (C3100, Intan Technologies).

### Two-photon calcium imaging

Imaging experiments started after mice were habituated for at least seven sessions to lick for the five gustatory stimuli. Images were acquired using a movable objective microscope (MOM, Sutter) with a resonant scanning module controlled by MScan (Sutter). The light source was a Ti:sapphire laser (Coherent) and an Olympus LCPLN20XIR objective (NA: 0.45, air, working distance: 8 mm) was used. GCaMP6f was excited at 940 nm with a laser power of 50-80mW at the front of the objective. Images (512 x 512 pixels) were acquired at 31 Hz with a 450 x 450 μm field of view (100-230 μm below the brain surface). Three hundred images (9.67 s long) were acquired for each trial (from 2 s before the cue to 7.67 s after the cue onset) and frame signals were synchronized with behavioral events through a RHD2000 recording system (Intan Technologies). In total, imaging data for 50-60 trials were acquired (10-12 trials for each tastant).

### Widefield calcium imaging

Images were acquired using a CMOS camera (FL3-U3-13E4M-C, FLIR) installed on the movable objective microscope (MOM, Sutter). The light source was a xenon arc lamp (Lambda LS, Sutter) filtered through a GFP filter cube. An Olympus XLFluor4x/340 objective (NA: 0.28, air, working distance: 29.5 mm) was used. FlyCapture (FLIR) was used to control imaging parameters and frames were taken at 16.6 Hz with a resolution of 1280 x 1024 pixels (~2 x 1.6 mm). Frame signals were synchronized with behavioral events through a RHD2000 recording system (Intan Technologies).

### Data Analysis

Data analysis was performed using ImageJ (NIH) and custom scripts written in MATLAB (MathWorks, Natick, MA).

### Behavioral analysis

The analog trace from the infrared beam was used for analyzing licking behaviors. A licking event was detected whenever the trace crossed a fixed threshold. Only licking within 7.5 s after the auditory cue were used for analysis (i.e., licking for rinses was not analyzed). A licking bout was defined as a train of at least three consecutive licks with an inter-lick interval shorter than 500 ms [39].

### Calcium imaging data analysis

For two-photon calcium imaging, images recorded for each trial (300 frames) were down-sampled from 31 Hz to 6.2 Hz with ImageJ (group z-projection) and concatenated across trials. Motion correction was performed with a package for piecewise rigid motion correction of calcium imaging data (NoRMCorre, https://github.com/flatironinstitute/NoRMCorre) [40]. Regions of interest (ROIs) corresponding to cell bodies, calcium traces and deconvolved activity were automatically extracted with the constrained nonnegative matrix factorization (CNMF)-based algorithm (https://github.com/flatironinstitute/CaImAn-MATLAB) [19]. The automatically detected ROIs were further manually corrected based on ROIs’ shape and calcium traces. For widefield imaging, videos were down-sampled from 16.6 Hz to 8.3 Hz with ImageJ (group z-projection). Motion correction was also performed with the NoRMCorre package. ROIs, calcium traces and deconvolved activity were automatically extracted with the CNMF-E package, an extension of the CNMF algorithm for one-photon imaging data (https://github.com/zhoupc/CNMF_E) [41]. The automatically detected ROIs were further manually corrected based on ROIs’ shape and calcium traces.

ROIs (putative cells) were categorized as responsive to cue, licking initiation or gustatory stimuli based on the deconvolved activity. Only excitatory responses were analyzed. For cue response, we compared the mean baseline activity (1 s before auditory tone) to the mean activity during stimulus, but before licking initiation (Wilcoxon rank sum test, p<0.05). For licking response, we compared the mean baseline activity (1 s before auditory tone) to the mean activity following the onset of the first lick, but before taste delivery, or 1 s following the first lick if taste delivery came later than 1 s (Wilcoxon rank sum test p<0.05). For cells that were responsive to both cue and licking, we further compared the mean activity before licking (0.5 s before licking) to the mean activity following licking (1 s or before taste delivery, Wilcoxon rank sum test p<0.05). This comparison was used to recategorize cells where observed licking response might be a carryover of the cue response. For taste responses, we compared the mean baseline activity (1 s before auditory tone) to the mean activity (1 s) centered on the peak of the response generated following taste delivery (within 3.5 s, Wilcoxon rank sum test, p<0.05). For cells responsive to all five gustatory stimuli and licking, we additionally compared the mean activity before taste delivery (0.5 s before taste delivery) to the mean activity (1 s) centered on the peak of the response generated following taste delivery (within 3.5 s, Wilcoxon rank sum test, p<0.05). This additional test was used to eliminate cells where observed gustatory response might be a carryover of the licking response. Taste-responsive neurons were also categorized based on their best responses. Specifically, a neuron’s best response was defined as the strongest taste-evoked response following taste delivery (within a 3.5 s window).

For hierarchical clustering analysis, taste-evoked deconvolved activity (peak response within 3.5 s after taste delivery) for each neuron was normalized to the maximum taste-evoked response (best response) for that cell. MATLAB functions including “linkage”, “cluster” and “dendrogram” were used to perform the agglomerative hierarchical clustering. Results of hierarchical clustering were also confirmed by using the evoked change of fluorescence intensity (ΔF/F).

To assess breadth of tuning [24], we calculated the entropy (H) for each taste-responsive neuron with the following equation: 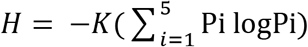, where Pi represents the proportional response to each of the 5 gustatory stimuli and K is a scaling constant (K = 1.431 for 5 tastants). Entropy value (H) ranges between 0 and 1, where 0 represents a neuron responds exclusive to one stimulus (narrowly tuned) and 1 presents a neuron responds equivalently to all 5 stimuli (broadly tuned).

To evaluate whether responses to taste qualities, cue and licking were spatially clustered, we calculated the pairwise distance between neurons (sessions with at least 3 neurons) responding to each of the five gustatory stimuli (16 out of 16 sessions), cue (15 out of 16 sessions) or licks (15 out of 16 sessions). We then calculated distance between the same number of randomly chosen neurons in the fields and repeated this procedure 1000 times. The mean distance between taste-responsive neurons was compared to the mean distance between randomly chosen neurons for each session (permutation test). Significant difference was defined based on whether the average distance between taste-responsive neurons was below the lowest 5% of the distance between random neurons (One-tailed permutation test, p<0.05). We also repeated this analysis with a stricter threshold (One-tailed permutation test, p<0.01). With this more stringent criterion, the distance between taste-responsive neurons was significantly smaller than the distance between randomly selected neurons in even fewer sessions than with the p<0.05 criterion. For the two-photon imaging dataset, we observed only three instances of significance (S:15/16 sessions, N: 16/16 sessions, CA: 16/16 sessions; Q: 15/16 sessions; W: 15/16 sessions), This was also the case for widefield imaging dataset, in which we observed only on instance where the distance between taste-responsive neurons was significantly smaller than the distance between randomly selected neurons (S:4 of 4 sessions, N: 4 of 4 sessions, CA: 4 of 4 sessions, Q: 4 of 4 sessions, W: 3 of 4 sessions).

To further quantify spatial clustering, we identified the centroids of clusters of neurons responding to each gustatory stimulus and compared intra-cluster distance with the inter-cluster distance. For instance, for neurons responding to sucrose, the intra-cluster distance was calculated as the distance between each neuron and the centroid of all neurons responding to sucrose (D_S-S_), the inter-cluster distance was calculated as the distance between each neuron responding to sucrose to the centroids of clusters of neurons responding to NaCl (D_S-N_), citric acid (D_S-CA_), quinine (D_S-Q_) and water (D_s-W_) (see **Supplementary Figure 1B** and **3C**). The normalized intra-cluster distance of neurons responding to each taste quality was calculated as the ratio between the intra-cluster distance and the average inter-cluster distance (**Figure 5E**).The intra-cluster and inter-cluster distance of neurons responding to cue, or lick was calculated in a similar way (**Supplementary Figure 2F**).

### Histological staining

Mice were deeply anesthetized with an intraperitoneal injection of a mixture of ketamine (140 mg/kg) and dexmedetomidine (2 mg/kg). Once fully anesthetized, mice were first intracardially perfused with 1x PBS followed by 4% paraformaldehyde. The brain was post-fixed overnight in 4% paraformaldehyde, then transferred to 30% sucrose until sunk (2-3 days). Brains were cut on a cryostat (HM505, Leica) into 50 μm coronal slices. Sections were washed in PBS, counterstained with Hoechst 33342 (1:5000 dilution, H3570, ThermoFisher, Waltham, MA), mounted on glass slides and imaged on a confocal microscope (LSM800, Zeiss).

---

**Supplementary Video 1: A video of neural activity from mouse gustatory cortex imaged through a prism with two-photo microscopy.** The video is played at 5 times of the actual speed.

**Supplementary Video 2: A video of neural activity from mouse gustatory cortex imaged through a prism with widefield imaging.** The video is played at 5 times of the actual speed.

