## supplementary figures for "Spatially distributed representation of taste quality in the gustatory insular cortex of awake behaving mice"

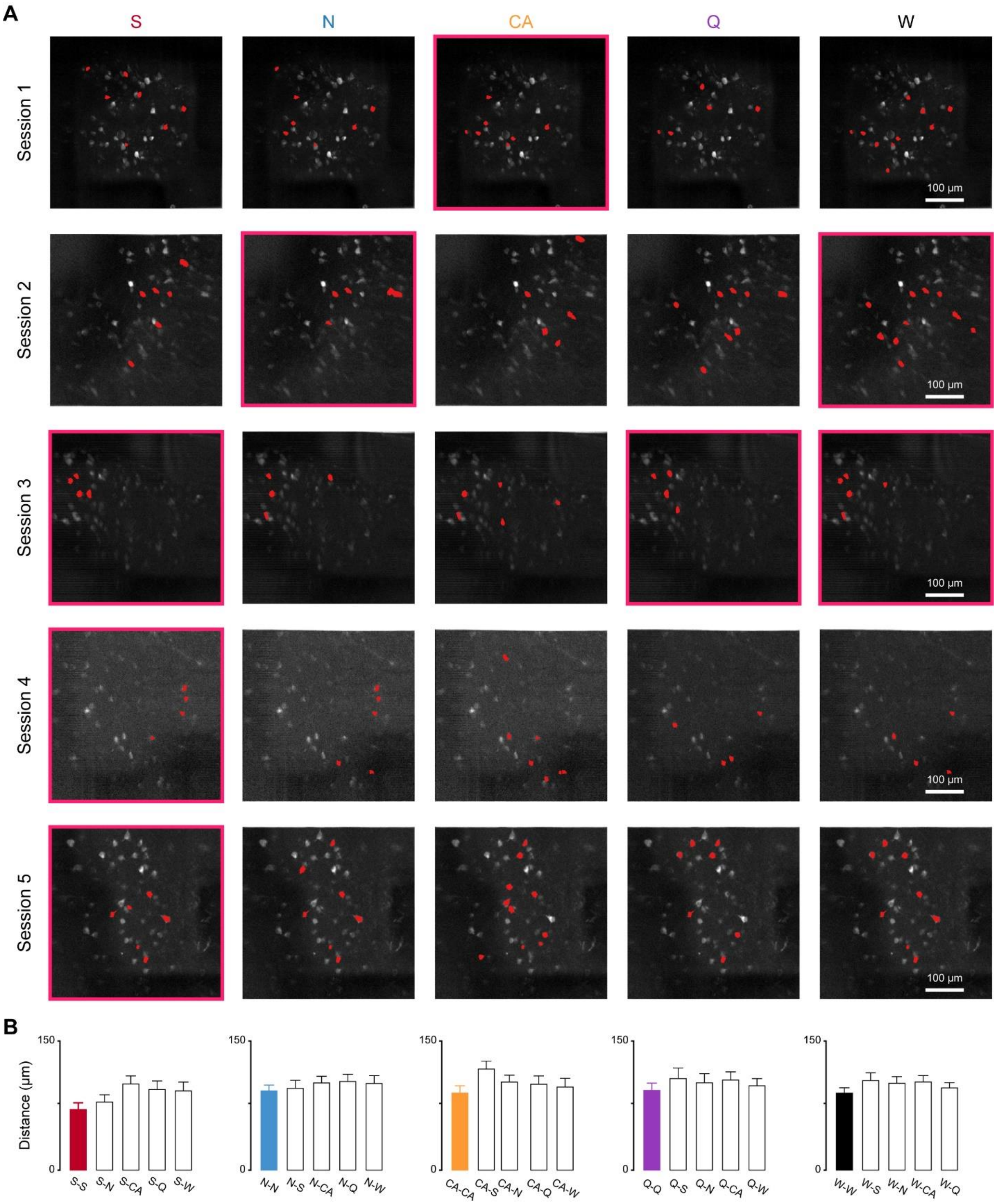


**Supplementary Figure 1: Intermingled representation of taste quality with two-photon imaging. A.** Two-photon images from 5 imaging sessions (4 mice) showing the location of neurons (red markers) responding to S, N, CA, Q and W. Each row represents an imaging session.

The images with red box represent stimulus representations in which the distance between taste-responsive neurons is significantly smaller than distance between random neurons ($p<0.05$). Notice that even in these sessions, responsive neurons for each taste quality were intermingled and on average 54% of neurons per session responding to multiple tastants. **B**. Bar graph showing the average intra-cluster distance and inter-cluster distance of neurons responding to each of the 5 gustatory stimuli (see methods). Here, only neurons in sessions shown in panel **A** were included. One-way ANOVA test, $p > 0.05$. Error bars represent SEM.

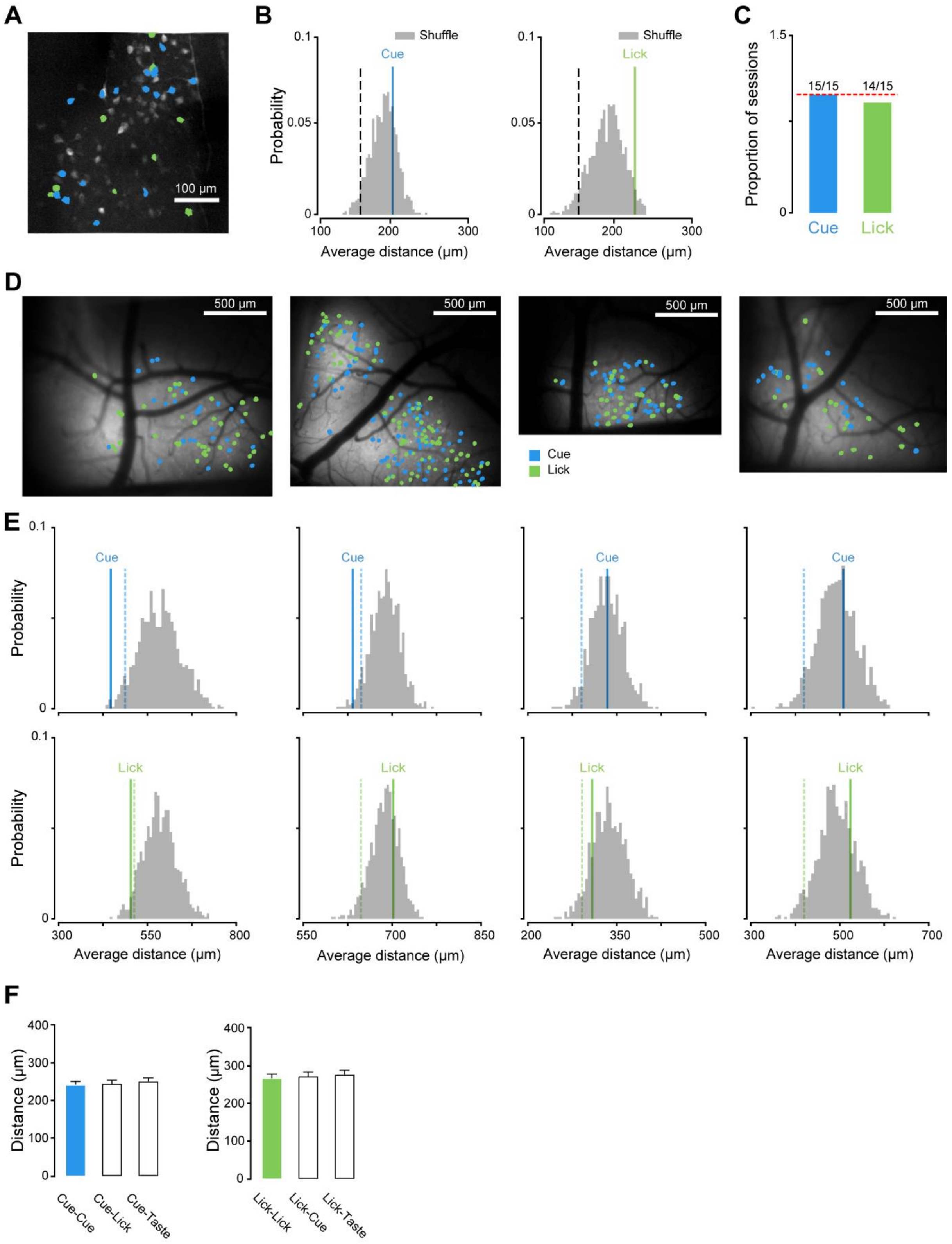


**Supplementary Figure 2: Intermingled representation of cue and licking. A.** A two-photon image showing the location of neurons responding to cue (blue) and lick (green). **B.**

Representative histograms showing the distribution of average pairwise distance between randomly chosen neurons (grey). The black dashed lines mark the boundary of the lowest 5% average pairwise distance for the random distribution. Colored lines represent the average pairwise distance between neurons responding to cue (blue) and lick (green), as shown in **A**. **C**. Bar plot showing the summary of imaging sessions in which distance between neurons responding to cue (15 out of15 sessions) and lick (14 out of15 sessions) were not significant different from null distribution. The red dashed line represents the 100%. **D**. Widefield images from 4 different sessions in 4 mice showing the location of neurons responding to cue (blue) and lick (green). **E.** Histograms showing the distribution of average pairwise distance between randomly chosen neurons (grey) from 4 sessions as shown in **D**. Each column represents a session. The black dashed lines mark the boundary of the lowest 5% average pairwise distance for the random distribution. Colored lines represent the average pairwise distance between neurons responding to cue (blue, top row) and lick (green, bottom row). **F**. Bar graph showing the average intra-cluster distance and inter-cluster distance of neurons responding to cue (left) and lick (see methods). Here, only neurons from the widefield imaging sessions shown in panel **D** were included. One-way ANOVA test, $p > 0.05$. Error bars represent SEM.

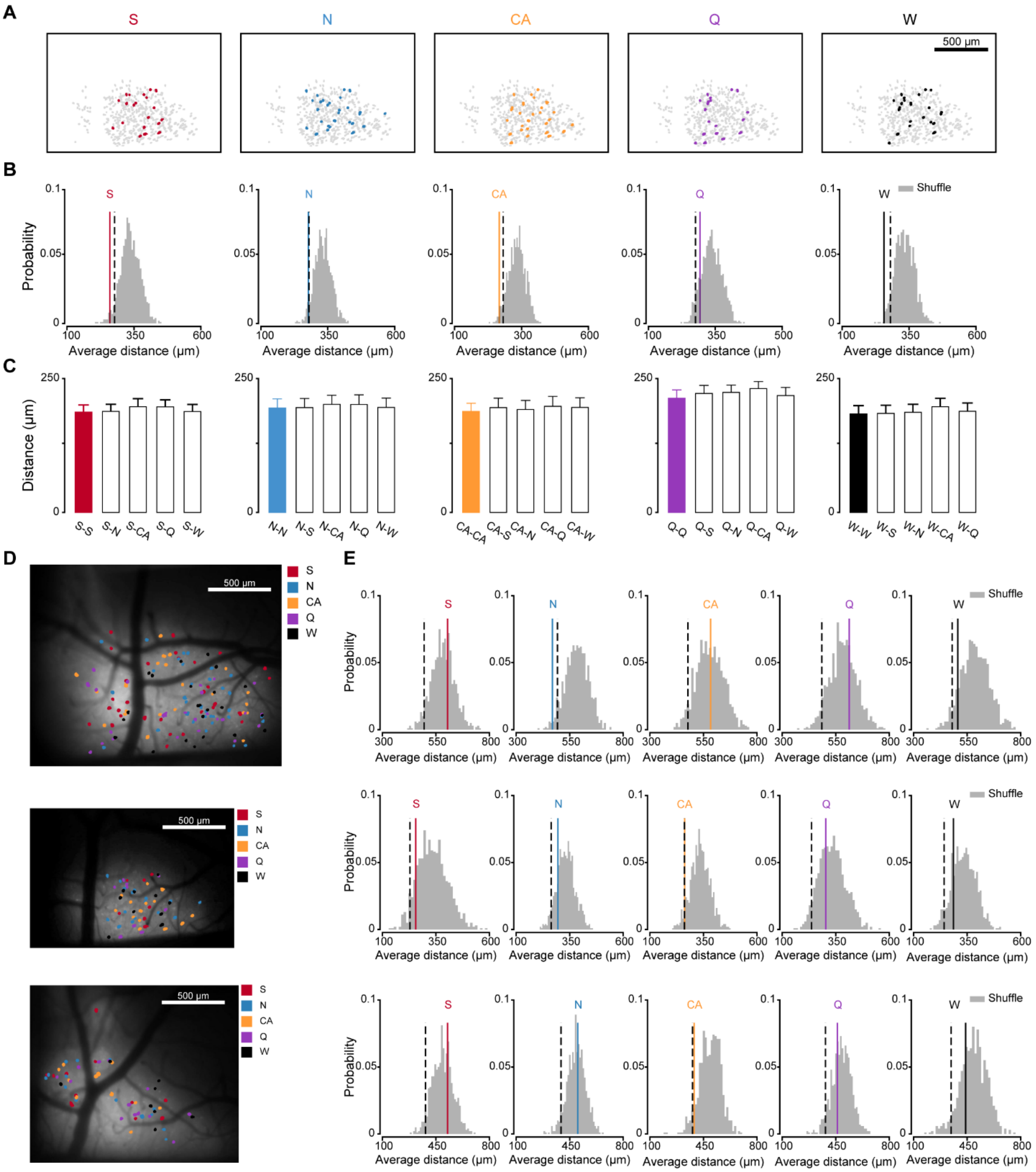


**Supplementary Figure 3: Intermingled representation of taste qualities with widefield imaging. A,** Spatial locations of neurons responding to S, N, CA, Q and W from an imaging session where distance between taste-responsive neurons was significantly smaller than null distribution ($p < 0.05$). Notice the intermingled organization of neurons responding to the four tastants. **B.** Histograms showing the distribution of average pairwise distance between randomly chosen neurons (grey). The black dashed lines mark the boundary of the lowest 5% average pairwise distance for the random distribution. Colored lines represent the average pairwise distance between neurons responding to S (red), N (blue), CA (orange), Q (purple) and W (black), as shown in **A. C.** Bar graph showing the average intra-cluster distance and inter-cluster distance of neurons responding to each of the five gustatory stimuli. Here, only neurons in the session shown in panel **A** were included. One-way ANOVA test, $p > 0.05$. Error bars represent SEM. **D.** Spatial map of neurons with best responses to S (red), N (blue), CA (orange), Q (purple) and W (black) from 3 different imaging sessions. **E**, Histograms showing the distribution of average pairwise distance between randomly chosen neurons (grey) from 3 different sessions as shown in **D**. Each row represents a session. The black dashed lines mark the boundary of the lowest 5% average pairwise distance for the random distribution. Colored lines represent the average pairwise distance between neurons with best response to S (red), N (blue), CA (orange), Q (purple) and W (black).
